# PURC v2.0: A Program for Improved Sequence Inference for Polyploid Phylogenetics and Other Manifestations of the Multiple-Copy Problem

**DOI:** 10.1101/2021.11.18.468666

**Authors:** Peter Schafran, Fay-Wei Li, Carl J. Rothfels

## Abstract

Inferring the true biological sequences from amplicon mixtures remains a difficult bioinformatic problem. The traditional approach is to cluster sequencing reads by similarity thresholds and treat the consensus sequence of each cluster as an “operational taxonomic unit” (OTU). Recently, this approach has been improved upon by model-based methods that correct PCR and sequencing errors in order to infer “amplicon sequence variants” (ASVs). To date, ASV approaches have been used primarily in metagenomics, but they are also useful for identifying allelic or paralogous variants and for determining homeologs in polyploid organisms. To facilitate the usage of ASV methods among polyploidy researchers, we incorporated ASV inference alongside OTU clustering in `PURC v2.0`, a major update to PURC (Pipeline for Untangling Reticulate Complexes). In addition to preserving original PURC functions, `PURC v2.0` allows users to process PacBio CCS/HiFi reads through DADA2 to generate and annotate ASVs for multiplexed data, with outputs including separate alignments for each locus ready for phylogenetic inference. In addition, `PURC v2.0` features faster demultiplexing than the original version and has been updated to be compatible with Python 3. In this chapter we present results indicating that `PURC v2.0` (using the ASV approach) is more likely to infer the correct biological sequences in comparison to the earlier OTU-based PURC, and describe how to prepare sequencing data, run `PURC v2.0` under several different modes, and interpret the output. We expect that `PURC v2.0` will provide biologists with a method for generating multi-locus “moderate data” datasets that are large enough to be phylogenetically informative and small enough for manual curation.

## 1 Overview

### 1.1 The Multiple-Copy Problem

There are many situations in biology in which an individual organism may harbor multiple closely related gene copies, and where knowing the number of copies and their individual sequences is important for downstream inferences. For example, a major system of self-incompatibility in plants (*e.g.*, the ability of a plant to reject fertilization attempts from its own pollen and thus avoid mating with itself) is based on the presence of particular combinations of “S-alleles”, and the characterization of these highly polymorphic S-alleles across individuals is thus essential for understanding the mechanism and consequences of this form of self-incompatibility (Ramanauskas and Igić, 2017; Goldberg et al., 2010). Similarly, many applications based on the multi-species coalescent, such as delimiting species with BPP (Yang and Rannala, 2010; Yang, 2015), rely on including the sequences from both alleles present in a diploid individual. This “multiple-copy problem” (Griffin et al., 2011), however, is particularly pronounced in phylogenetic studies of polyploids. Because many polyploids unite subgenomes from different progenitor species (they are hybrid polyploids, or “allopolyploids”), their true evolutionary history is a network rather than a strictly bifurcating tree, and to infer this reticulate history the homeologous sequences—the copies from each of the subgenomes—need to be recovered and reconstructed accurately (Rothfels, 2021).

The classic way of recovering multiple gene copies from a single individual is molecular cloning: the desired marker is amplified by PCR, the amplicon is cloned into plasmid vectors that are used to transform *Escherichia coli*, and multiple colonies of the transformed bacteria are re-amplified and sequenced (*e.g.*, Schuettpelz et al., 2008; Li et al., 2012). This approach is labor-intensive and expensive (for a triploid, for example, one would need to sequence 11 colonies to have a 95% chance of getting all three copies), making datasets with many samples or many loci impractical. Short-read next-generation sequencers, such as those of the Illumina platform, offer some relief, in that the reads come from individual molecules (rather than representing a form of majority-rule consensus of the molecules present, as is the case with Sanger sequencing) and sequencing costs are dramatically reduced. However, in order to recover the individual copies, the reads need to be assembled accurately. This assembly step can be bioinformatically prohibitive, especially if there are more than two copies present, and always faces a hard limit: to correctly assemble the full length of each copy of a target sequence, consecutive variable sites have to be separated from each other by no longer than the read length.

### 1.2 The PURC Approach

To help facilitate the recovery of all homeologous sequences from polyploid accessions (and for other manifestations of the multiple-copy problem), Rothfels et al. (2017) published a molecular lab workflow based on PacBio long-read sequencing, with an associated bioinformatic pipeline (the “Pipeline for Untangling Reticulate Complexes”, PURC) for inferring the individual copies from the PacBio reads. This approach capitalized on PacBio’s circular consensus sequencing (CCS) technology to generate contiguous reads for long (*>*1000 bp) phylogenetically informative regions, thus avoiding the need for assembly and allowing for the accurate retrieval of all copies present in individual polyploids (Rothfels et al., 2017; Dauphin et al., 2018; Kao et al., 2020). A single PacBio SMRT cell can generate sequences for multiple loci for hundreds of samples, providing an economical means to generate powerful “moderate data” datasets for polyploid phylogenetics.

The wetlab workflow involved standard PCR with barcoded forward primers; the amplicons were then pooled (roughly standardized to equal concentration taking into account the anticipated number of copies present in each accession) and sequenced on a single PacBio SMRT cell. PURC can demultiplex the resulting reads by locus, barcode, and phylogenetic affinity (*i.e.*, the same barcode can be used multiple times in a single sequencing run, once for each phylogenetically-discernable group, such as a genus). By taking an iterative chimera-removal and operational taxonomic unit (OTU) clustering approach (using USEARCH and UCHIME; Edgar, 2010; Edgar et al., 2011), PURC then infers the underlying biological sequences. In brief, OTU clustering infers groups of reads that fall within a set level of similarity to the “centroid” read representing the middle of a group, and creates a single consensus sequence from all reads within each group. In this context, it’s assumed that each group would represent a unique homeologous copy. The output from PURC is one alignment for each locus, which includes all the copies present in each of the accessions, labelled by their accession ID and coverage (the number of reads that constitute that sequence). Since its release, PURC has been used to investigate allopolyploidy in genus-level datasets in diverse plant lineages (Morales-Briones and Tank, 2019; Dauphin et al., 2018; Suissa et al., 2020; Blischak et al., 2020), in studies of cytotype variation within species (Kao et al., 2020), and for applications where polyploidy itself was incidental to the primary research questions (Kao et al., 2019; Chery et al., 2019; Wolfe et al., 2021; Frost et al., 2020). Most notably, Blischak et al. (2018) created a program to adapt PURC to data generated by microfluidic PCR, reducing one of the main limitations of amplicon-based approaches (the time and expense associated with PCR itself).

### 1.3 PURC v2.0

Since the release of `PURC`, several studies have demonstrated shortcomings of OTU clustering, which is the method `PURC` used to infer biological sequences. In particular, OTU clustering tends to over-estimate the number of sequences, is sometime unreproducible (repeat runs yield different OTUs), and it is difficult to determine appropriate similarity thresholds (Callahan et al., 2017; Barnes et al., 2020; Joos et al., 2020; Nelson et al., 2020), with the overestimation problem reported for `PURC` specifically (Morales-Briones and Tank, 2019; Blischak et al., 2018). An alternative approach to identify and separate PCR and sequencing errors from biological sequences is to apply an error model, where read abundance, composition, and quality scores are used to infer whether each unique read is likely to have been derived from another observed sequence (Callahan et al., 2016). Reads inferred to represent biological sequences by these methods are called amplicon sequence variants (ASVs), exact sequence variants, or zero-radius OTUs. `DADA2` (Callahan et al., 2016) is one of the most popular software packages for inferring ASVs, and is particularly flexible because it incorporates separate error models for Illumina and PacBio CCS data. Additionally, `DADA2` can take sequences as “priors”, increasing the sensitivity of the algorithm for sequences that are similar to the priors (see description under `dada` function in the `DADA2` version 1.12.0 manual and detailed usage at https://benjjneb.github.io/dada2/pseudo.html).

In order to take advantage of the potential improvements offered by ASVs, `PURC v2.0` incorporates `DADA2` alongside `Vsearch` (Rognes et al., 2016), an open-source alternative to `Usearch` (Edgar, 2010), allowing `PURC v2.0` to run in four different ways:

1. OTU clustering alone
2. ASV inference alone
3. OTU clustering and ASV inference in one run
4. OTU clustering followed by ASV inference with the OTUs as priors

To test our incorporation of `DADA2` we reproduced OTUs and ASVs generated by Nelson et al. (2020), where a mock community of five cyanobacteria was created by combining equal quantities of *rbcL-X* amplicons generated from pure cultures. `PURC v2.0` was run on four replicates of the five-taxon mock community, performing OTU clustering with default parameter values (clustering thresholds by round: 0.997, 0.995, 0.990, 0.997, final size threshold = 4) and inferring ASVs (`DADA2`) with a maximum of five expected errors allowed per read and length outliers removed based on Tukey’s outlier test (Tukey, 1977). We also tested data from species of *Isoetes* from Suissa et al. (2020) using the same parameters.

Our results agree with Nelson et al. (2020) in showing that ASV inference was more accurate than OTU clustering (Figure 1). The same five ASVs were recovered from every replicate, whereas OTU clustering identified six to nine OTUs per replicate. Most replicates contained OTUs identical to ASVs, but OTUs with small sequence deviations were common and a few spurious OTUs with 75%–92% identity to ASVs were also generated. Because the *Isoetes* data from Suissa et al. (2020) are empirical and the true sequences are unknown, we evaluated ASV-versus OTU-based inferences of these sequences using two independent PCR amplifications of each of three individuals, thought to represent one diploid and two allotetraploids (*I. echinospora* Taylor #6989-1, *I. maritima* Taylor #6983, and *I.* aff. “new hybrid A” Taylor #6988-2, respectively). For these accessions we inferred 20 OTUs versus 10 ASVs in total. Both PCR replicates yielded identical ASV inferences: one sequence for the diploid, and two for each of the allotetraploids (Figure 2) and these sequences all had high coverage (they represented many individual reads; Figure 3). However, OTU inference was less consistent. For the diploid, both PCR replicates resulted in a high-coverage OTU that was identical to the one from ASV inference, but one of the replicates inferred two additional low-coverage OTUs (for example, OTU-6 contained five deletions and two ambiguous nucleotides relative to OTU-412 and ASV-310; Figure 2A.) Similarly, for the tetraploids, for both PCR replicates, OTU inference found the sequences corresponding to the ASVs, but also found additional, presumably spurious, low-coverage sequences (Figures 2, 3). One of these sequences—OTU-5 in replicate 2 of the allotetraploid *I.* aff. “new hybrid A” Taylor #6988-2—was uniquely divergent, with 36 SNPs and two deletions totaling 97 bases. The origins of the five reads that constitute this OTU are unclear, but were not due to low-quality basecalls or incorrect assignment by barcodes. In every case, the identical ASVs and OTUs were by far the most abundant (Figure 3); they likely represent the true biological sequences.

**Figure 1:**
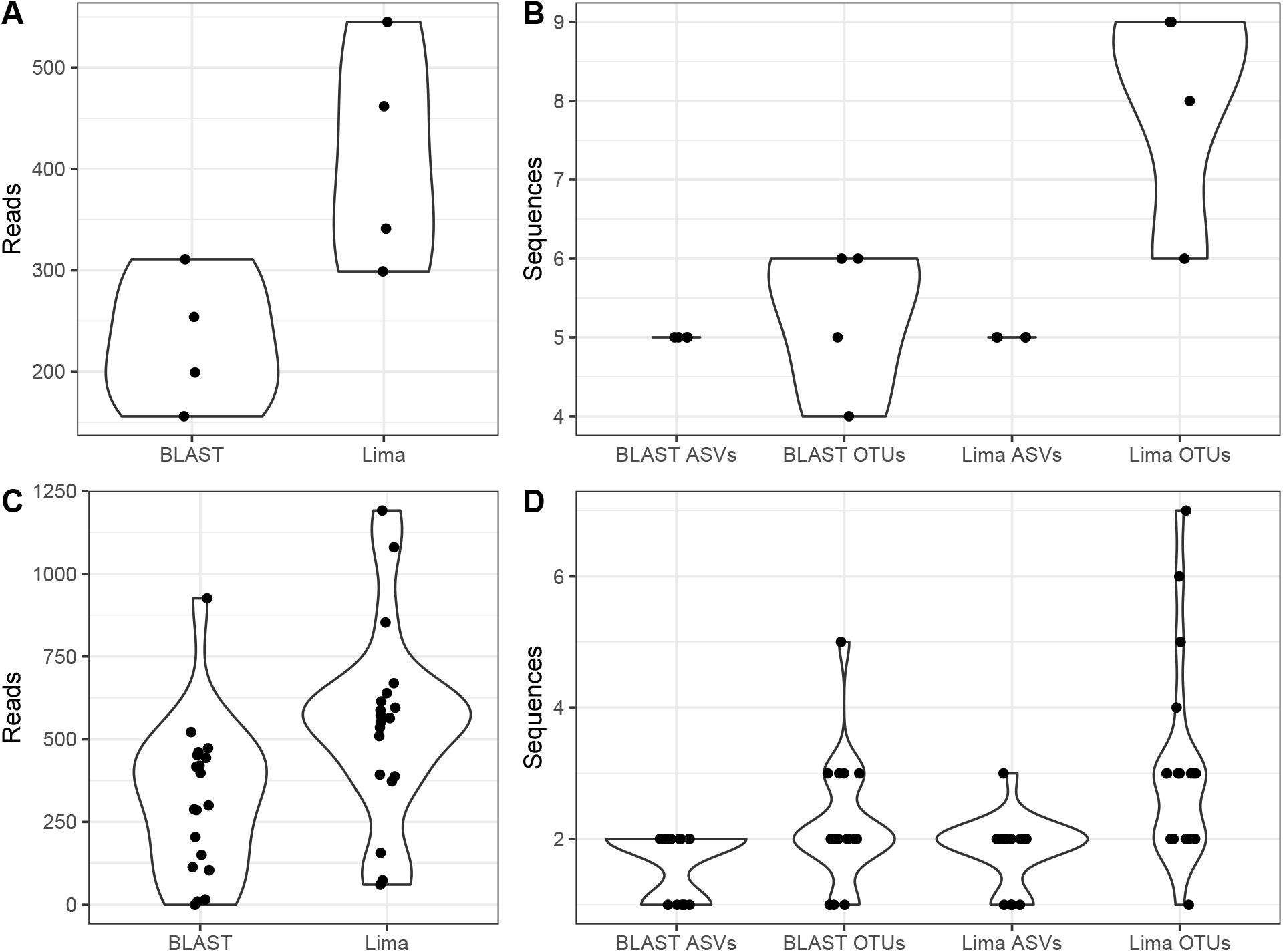
Comparison of demultiplexing and sequence inference methods using data from Nelson et al. (2020) and Suissa et al. (2020). *A*: Number of reads assigned to four replicates of the five taxon mock community. *B* : Sequences inferred for four replicates of the five taxon mock community. *C* : Number of reads assigned to each *Isoetes* sample. *D* : Sequences inferred for each *Isoetes* sample.

**Figure 2:**
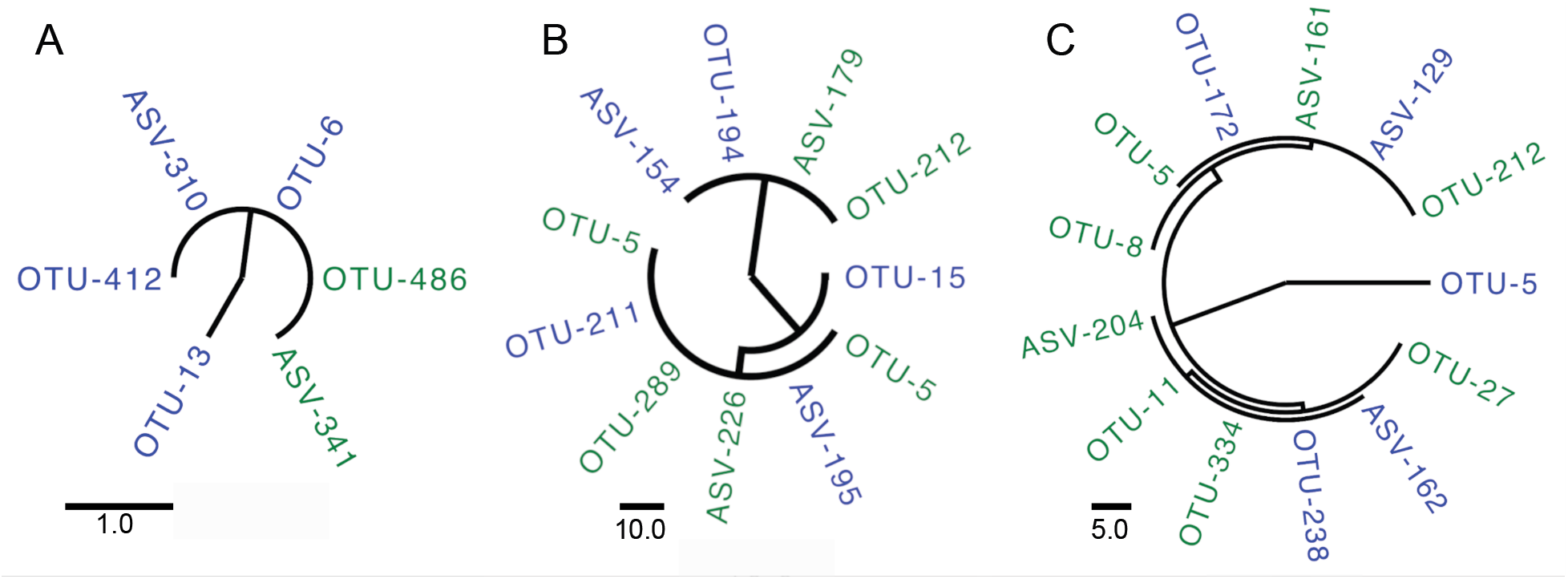
Maximum-parsimony trees of OTUs and ASVs from two PCR replicates of three *Isoetes* species (A: diploid *I. echinospora* Taylor #6989-1; B: allotetraploid *I. maritima* Taylor #6983; C: allotetraploid *I.* aff. “new hybrid A” Taylor #6988-2. PCR replicate 1 is colored green and replicate 2 is colored blue. Numbers following tip names indicate coverage (the number of reads constituting each OTU/ASV). Trees are rooted at their midpoints. Scale bar represents number of substitutions. Data are from Suissa et al. (2020).

**Figure 3:**
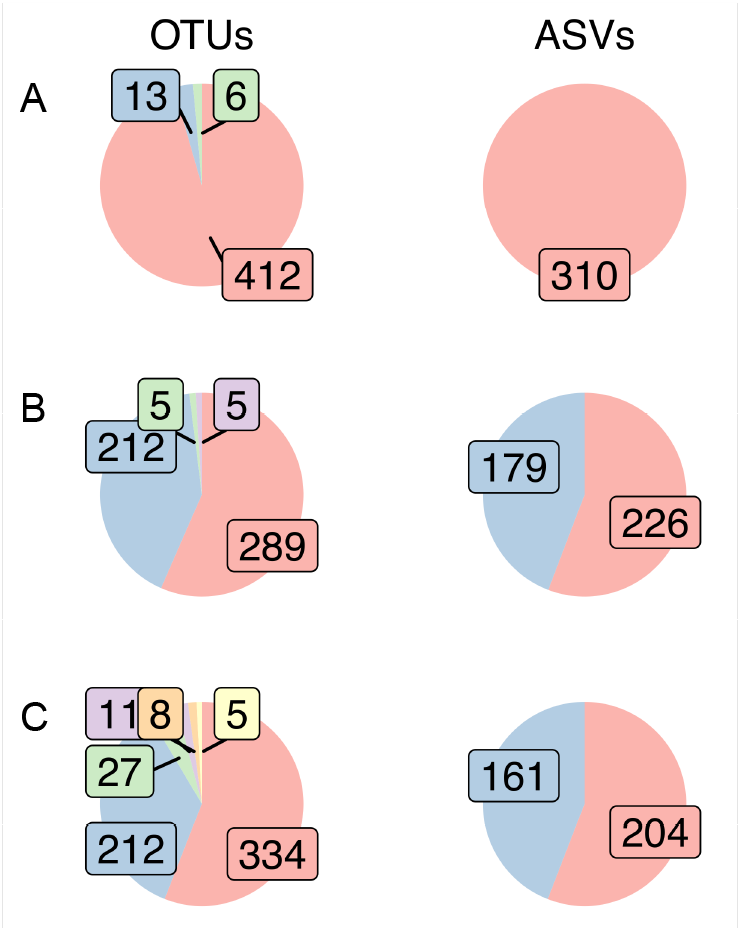
Proportions of reads contributing to each OTU and ASV for three *Isoetes* species (A: diploid *I. echinospora* Taylor #6989-1; B: allotetraploid *I. maritima* Taylor #6983; C: allotetraploid *I.* aff. “new hybrid A” Taylor #6988-2, from one PCR replicate. Labels indicate number of reads in each OTU/ASV and sequences are colored from largest to smallest within each chart. Data are from Suissa et al. (2020).

`PURC v2.0` also includes PacBio’s lima tool (https://lima.how) as a new method for demultiplexing reads on Linux operating systems. Using lima, we could recover an average of 74% more reads from the Suissa et al. (2020) data than with the original BLAST-based method in `PURC`, including recovering one individual for which no sequences were identified using BLAST; the per-sample increase ranged from 24–640% (Figure 1 C). The number of both OTUs and ASVs increased in the `lima`-demultiplexed dataset but ASVs fluctuated less, with 74% of samples having the same number of ASVs as in the BLAST-demultiplexed data versus only 26% for OTUs.

There were some consistent trends in our analyses of mock community and real polyploid data. lima consistently recovered more reads from each sample, which resulted in an increase in the number of OTUs, possibly by inclusion of more divergent reads rising above the threshold for dropping low-abundance clusters. However, the inclusion of the additional reads had a much smaller effect on ASV inference. The variance in number of ASVs was always smaller than that for OTUs, and especially in the mock community analyses where every replicate was correctly inferred (Figure 1). While Nelson et al. (2020) showed that by varying OTU clustering parameters they could more accurately reconstruct the true sample composition, this approach is unreliable in cases where the true sequences are unknown and where it may be tempting to change parameters until the results match expectations. Based on our results, we recommend `lima` demultiplexing and ASV inference as the primary method for running `PURC v2.0`. To compare OTU clustering and ASV inference on your own data, `PURC v2.0` allows generating both simultaneously, with summary files to examine number and abundance of sequences for each sample. If it appears that too few ASVs are found, a new analysis can be run with the OTU sequences as priors, increasing the sensitivity of the DADA2 algorithm and reducing the detection limit for variants. This approach may be particular useful for samples with little data or where two biological sequences are very similar, although it remains to be tested in polyploids.

## 2 Materials

### 2.1 Hardware

A personal computer with a multi-core processor and at least 8 GB RAM, though precise requirements will vary based on the input data size.

### 2.2 Software

1. A Unix- or Linux-based operating system (*e.g.*, macOS, Ubuntu); on Windows, a Linux virtual machine can be used.
2. Conda package manager (https://docs.conda.io)

## 3 Methods

The following instructions as well as additional information for troubleshooting can be found in the main `PURC v2.0` repository at https://bitbucket.org/peter_schafran/PURC/.

### 3.1 Installing PURC v2.0

The most up-to-date version of `PURC v2.0` can be downloaded as a compressed TAR file in the main repository. When uncompressed, a new directory named `purc` will contain executable and installation files.

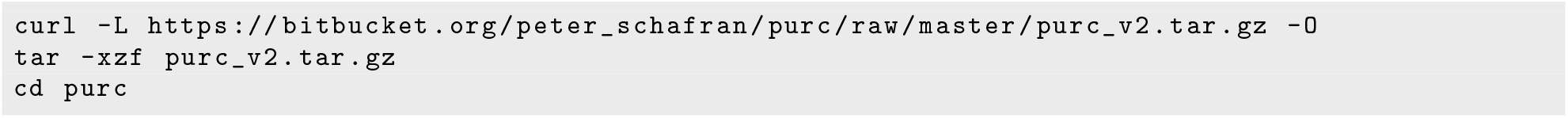

We recommend installing dependencies through Conda. Two YAML files, one for macOS and one for Linux, are included in the repository, and can be used to create a new `PURC v2.0` environment containing all necessary dependencies with one of these commands:

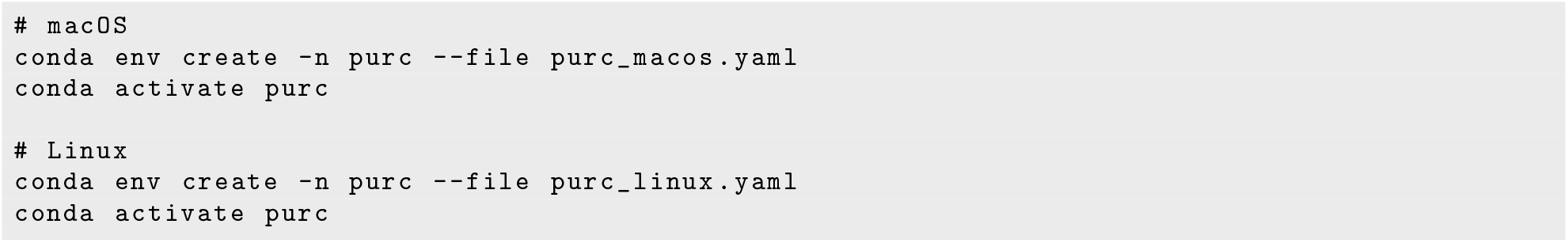

### 3.2 Preparing Input Files

Sequence data are expected to be *>*99% accurate amplicon sequences containing sample-specific nucleotide sequences (barcodes) on one or both ends of each read, as well as the priming sites used to generate the amplicons. If using PacBio reads generated by their standard protocol (https://ccs.how), they should not need any modification. Either FASTA or FASTQ formatted data are accepted, though FASTQ is required to perform ASV inference. In addition to the data, `PURC v2.0` requires four files:

1. Barcode file listing the barcode sequences
2. Reference sequences file, so that reads can be oriented, assigned to the correct locus, and sorted into phylogenetic groups if individual barcodes are used for multiple accessions
3. Map file(s), which link barcode and group IDs to unique accessions (one map file for each locus)
4. Configuration file, which includes information on specific settings, the primer sequences, and other necessary information

#### 3.2.1 Barcode File

Barcode sequences are provided in FASTA format. The sequence ID is the barcode name, and sequences should be oriented in the 5’-3’ direction. For example:

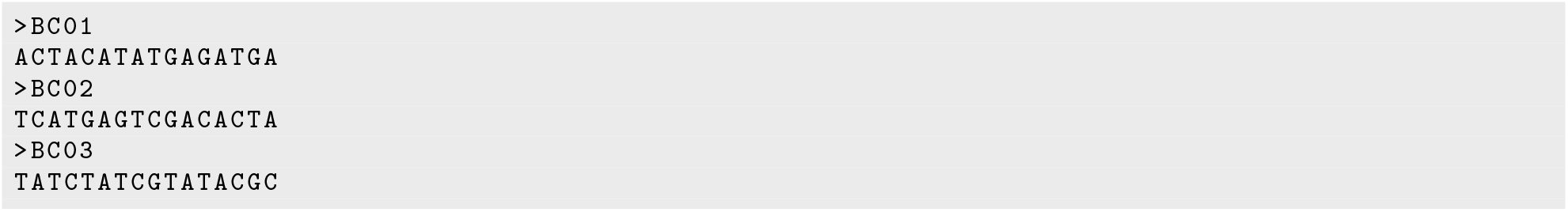

If using the dual barcode function, the barcode names must be BCF1, BCF2, . . . and BCR1, BCR2, . . . for barcodes on forward and reverse primers, respectively. This information is used to check for barcodes found in the incorrect orientation. We recommend barcodes that are unique (including reverse complements), especially if using the option to demultiplex with `lima`. However, if a subset of duplicate barcodes are detected, `PURC v2.0` will attempt to run separate analyses through `lima` with the unique and duplicate barcodes, and then merge the output.

#### 3.2.2 Reference Sequence File

Reference sequences must be provided in FASTA format. Each sequence should specify its locus name in the sequence ID (*e.g.*, locus=ApP), even if the data being analyzed represent only one locus. A group name (*e.g.*, group=A) can be provided to demultiplex by BLAST comparison to reference sequences if multiple samples share barcodes. A taxon name (*e.g.*, ref_taxon=Cystopteris bulbifera) or other descriptor can be included to provide additional information for the user but this information is not used by `PURC v2.0`. Fields are separated by forward slashes, so a complete sequence ID line may look like:

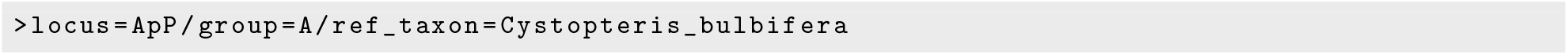

Note that the total sequence ID should not exceed 50 characters or else BLAST database construction will fail. All reference sequences for a locus must be in the same orientation to allow proper orientation of the reads.

#### 3.2.3 Map File

The map file indicates which barcodes and/or groups correspond with each sample. This file is a tab-delimited text file with one of three configurations:

1. If each sample contains just one barcode (*i.e.*, only the forward primer is barcoded) and each barcode is unique to one sample, the first column contains barcode names (from the barcode FASTA file) and the second column contains the name of the corresponding sample.

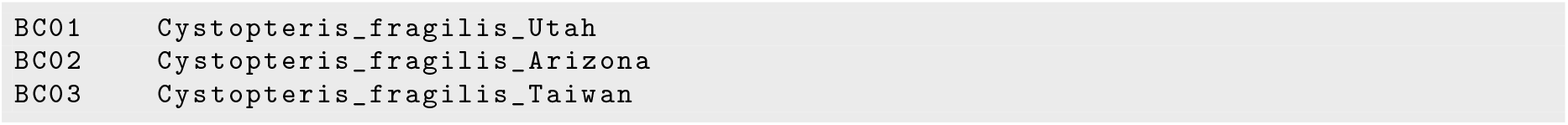
2. If each sample contains one barcode, but individual barcodes are used for multiple samples, then the first column contains the barcode name, the second column contains a group ID (corresponding to a group specified in a reference sequence ID), and the third column contains the sample name.

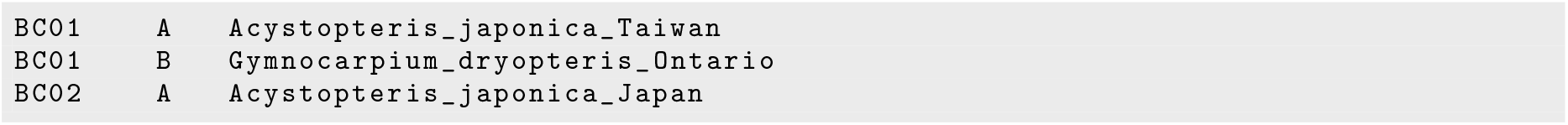
3. If samples are dual-barcoded, the first column contains the forward primer barcode name, the second column contains the reverse primer barcode name, and the third column the sample name.

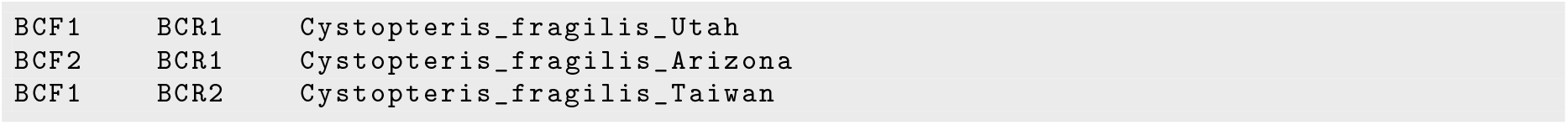

#### 3.2.4 Config File

All `PURC v2.0` operation is controlled via the config file, a text file containing information such as file names and run parameters. Paths to the data file, barcode file, reference sequence file, and map files are specified here, as well as primer sequences, run options, and optional parameter settings. Each line item is described by comments in the file (not shown here).

The first section of the config file specifies input files for reads, barcodes, and references, and the name of the prefix appended to output files, the output folder, and the log file.

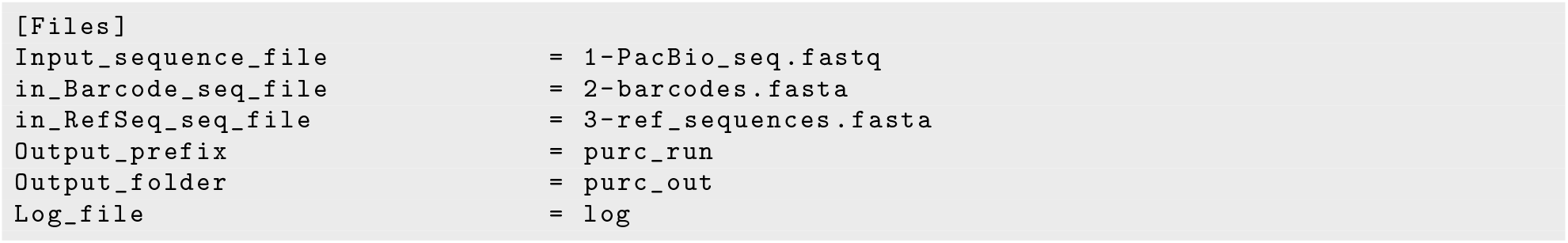

The second section provides information about the locus or loci to be processed. The locus name must match that provided in the reference sequence file. If processing multiple loci, their order in Locus name and `Locus-barcode-taxon_map` must match.

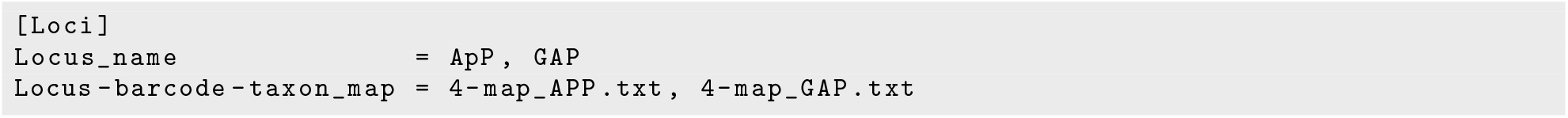

The third section is used to provide primer sequences in the 5’-3’ direction: these, too, must be in the same order as, *e.g.*, `Locus_name`. IUPAC ambiguous nucleotide codes are accepted.

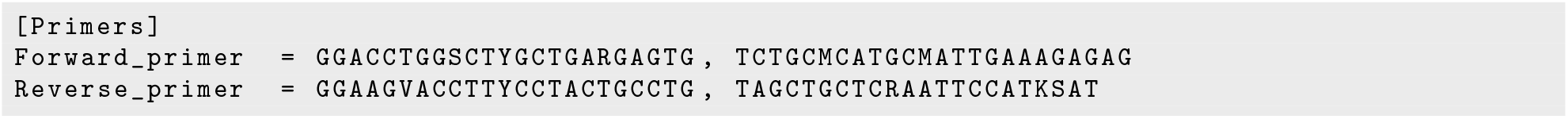

The fourth section allows the user to choose between several run modes depending on their data.

- Mode controls whether `PURC v2.0` checks for interlocus concatemers or not (`Mode` should be set to 1 for single-locus data).
- `Multiplex_per_barcode` indicates if barcodes are reused for multiple individuals within the same locus or if each barcode (for a given locus) is unique to one sample (barcode reuse is not available for dual barcoding).
- `Dual_barcode` is used to specify how the barcodes are arranged on each read. Note that options `1` and `2` are only treated differently if using `lima` to demultiplex.
- `Barcode_detection` describes where to look for barcodes in each read. If set to `0`, the “ErrMidBC” flag is included in the sequence name if barcodes are not at the ends of the sequence. This setting has no effect if using `lima`.
- `Recycle_chimeric_seq` controls whether `PURC v2.0` splits interlocus chimeras into their respective loci and includes them in downstream analysis, or discards such chimeras. This setting has no effect if `Mode = 1`.
- `Recycle_no_barcoded_seq` allows for the use of Smith-Waterman local alignment (Smith and Waterman, 1981) to try to identify barcodes in any sequences that failed to have a significant BLAST match. This setting has no effect if using `lima` to demultiplex.
- `Clustering_method` specifies which method to use for sequence inference. Option `2` does ASV inference and OTU clustering together, and is required for `Use_OTU_priors = TRUE`.
- `Align` controls whether each final sequence file produced (one per locus per clustering method) is aligned using `MAFFT` (Katoh and Standley, 2013).

For example:

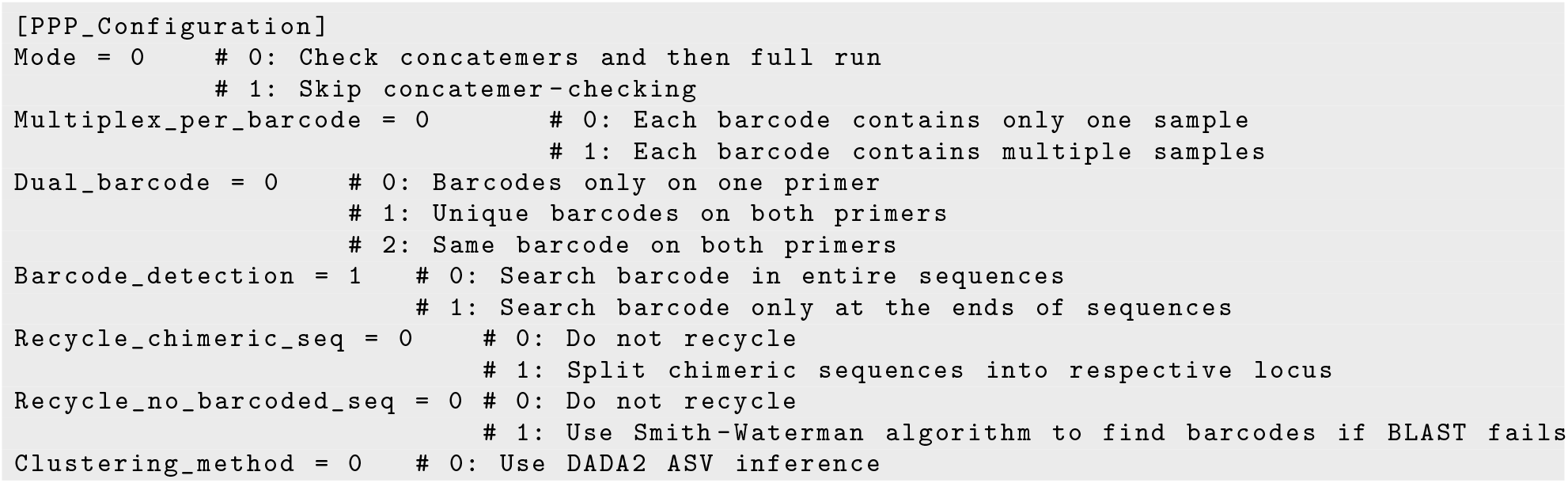

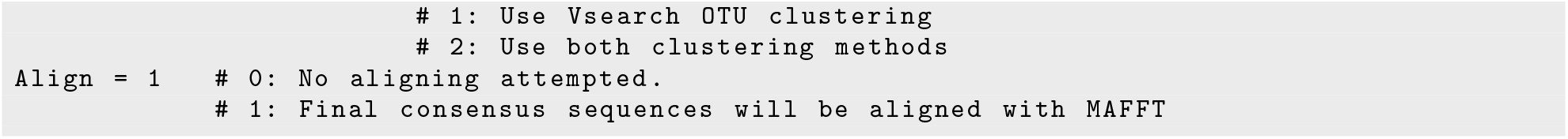

Other optional parameters follow this section of the config file, including those to change the operation of `DADA2` and `Vsearch. DADA2` options include specifying minimum and maximum lengths and maximum number of expected errors for reads to be included, and whether to include each sample’s OTU sequences as priors. If `minLen` and/or `maxLen` is set to 0, that parameter is calculated using Tukey’s equation for outliers (Tukey, 1977):

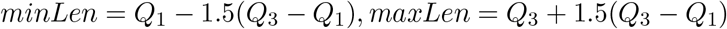

where *Q*_1_ is the first quartile and *Q*_3_ the third quartile, and results are rounded to the nearest integer. Outliers are recalculated for each sample. If `minLen` and/or `maxLen` values are user-supplied, they are applied globally. The maximum number of expected errors is estimated by `DADA2` based on read quality scores. If putative spurious ASVs are produced this number can be reduced, while if too many reads are discarded during filtering it can be increased.

- `minLen` is the minimum length for a read to be included in analysis. Set to 0 to automatically detect short outliers
- `maxLen` is the maximum length for a read to be included in analysis. Set to 0 to automatically detect long outliers
- `maxEE` is the maximum number of expected errors estimated by `DADA2` based on read quality scores. Reads with more expected errors than `maxEE` are discarded.
- `Use_OTU_priors` determines whether to use the OTU sequences as priors. When activated, the OTUs for each sample are used to infer ASVs for that sample, which can be useful if the number of reads per sample is low, or if well-supported OTUs seem to be lost during ASV inference. However, this option can increase the risk of creating spurious ASVs; it requires `Clustering_method = 2`.

For example:

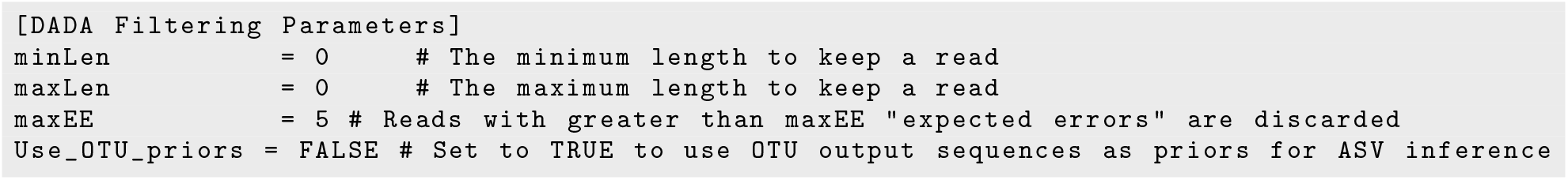

Available `Vsearch` options are for the identity levels for each round of clustering, minimum size to retain a cluster, and the abundance skew for detecting chimeric sequences.

- clustID*n* specifies the identity threshold for clustering reads during the *n*th round of clustering. For example, 0.997 means a read must have 99.7% identity to a cluster’s centroid sequence in order to be included in that cluster.
- `sizeThreshold1` sets the minimum number of reads in a cluster for it to be retained for the next round of OTU clustering.
- `sizeThreshold2` is the same as `sizeThreshold1`, but is only applied to the final clustering output.
- `abundance_skew` is the minimum ratio of parent sequences to putative chimeric sequence required to classify a sequence as chimeric. Parent sequences are expected to be at least twice as abundant as their chimeras.

For example:

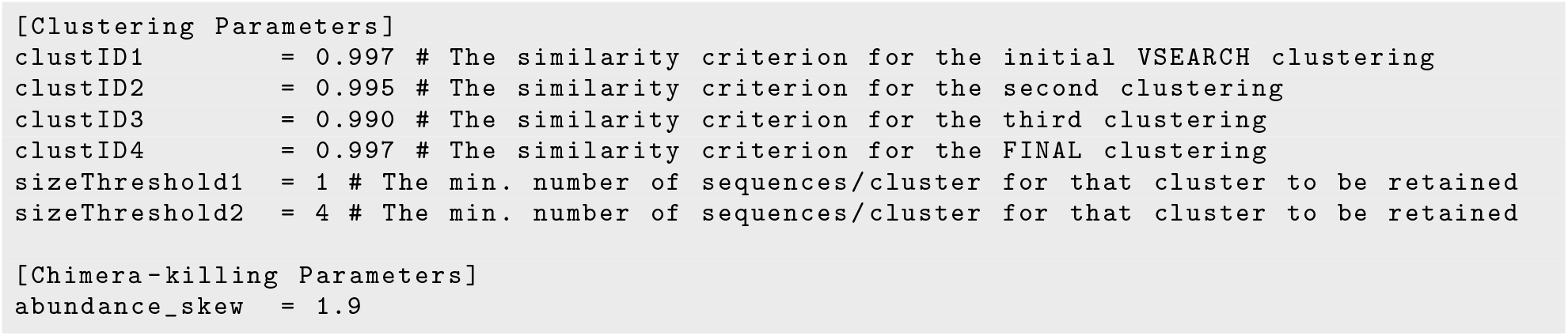

If the user’s operating system is detected as Linux-based, `PURC v2.0` will default to using `lima` for demultiplexing. To override and use BLAST-based demultiplexing, change the override flag in the config file.

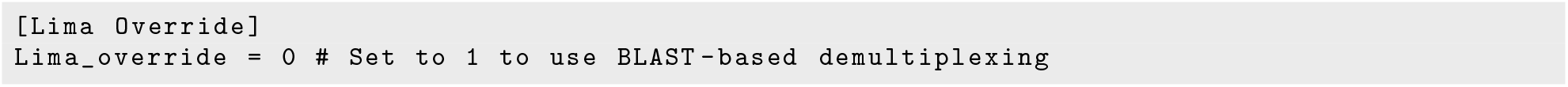

### 3.3 Running PURC v2.0

#### 3.3.1 Full Run With Demultiplexing and Sequence Inference

Once all input files are complete, `PURC v2.0` is initiated by calling the main `PURC` script with the config file as the argument:

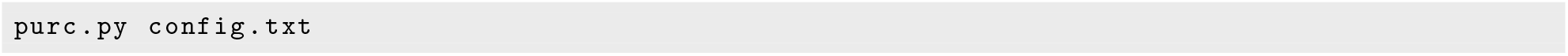

On completion, the output directory will have the following structure, where log, output prefix, locus, and clustering method are replaced with those specified by the config file. It will contain a subdirectory for each locus, and within each locus directory a subdirectory for each sample.

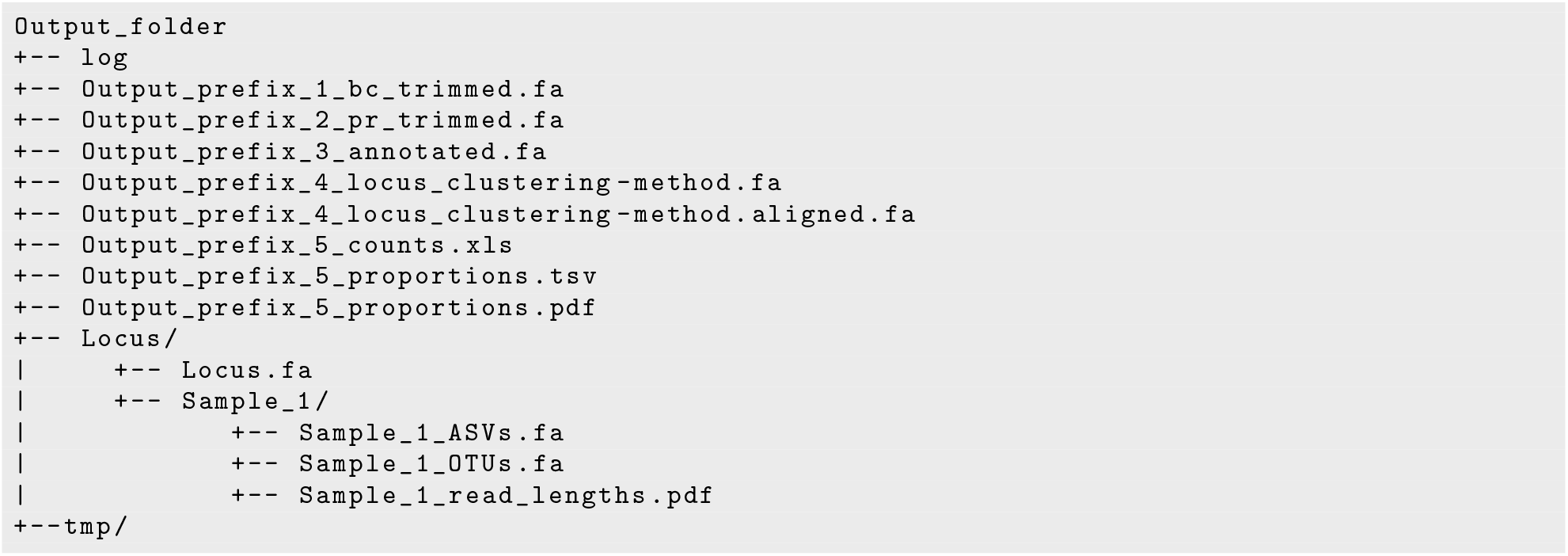

- log — log file documenting the `PURC` run
- Output_prefix_1_bc_trimmed.fa — all reads containing valid barcodes
- Output_prefix_2_pr_trimmed.fa — reads with primer sequences removed (OTU clustering only)
- Output_prefix_3_annotated.fa — reads that could be assigned to samples based on barcodes or groups
- Output_prefix_4_locus_clustering-method.fa — combined output sequences from all samples following OTU clustering and/or ASV inference. One file per locus per clustering method
- Output_prefix_4_locus_clustering-method.aligned.fa — alignment of output sequence file
- Output_prefix_5_counts.xls — summary of results containing number of reads surviving at each step in processing and the final number of sequences per sample
- Output_prefix_5_proportions(.tsv/.pdf) — summary of read coverage for the OTUs/ASVs for each sample. The PDF presents these data as pie charts with labels indicating the read coverage of each slice (as in Figure 3)
- Locus/Locus.fa — all reads annotated to this locus
- Locus/Sample_1/Sample_1_ASVs.fa — final ASV sequences for this sample (if applicable)
- Locus/Sample_1/Sample_1_OTUs.fa — final OTU sequences for this sample (if applicable)
- Locus/Sample_1/Sample_1_read_lengths.pdf — histogram of read lengths prior to ASV inference. Dotted lines indicate the limits for discarding too short/too long reads

If `PURC v2.0` is interrupted, it can be resumed by running again with the same config file. As long as all parameters are the same, it will determine the last completed step and continue.

#### 3.3.2 Analyses On Previously Demultiplexed Data

`PURC v2.0` includes a secondary script, `purc_recluster.py`, that can be used to perform OTU clustering on prior `PURC` runs or on data that have been demultiplexed by another method. Unlike the main script, `purc_recluster.py` operates with only command-line arguments.

Usage:

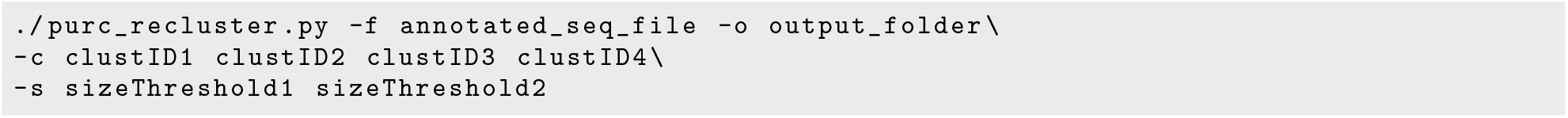

If using `PURC v2.0` demultiplexed data, the input file is the `output_prefix_3_annotated.fa` file. If preparing your own data, the file must be FASTA formatted with name lines structured as follows:

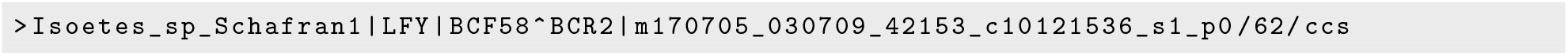

The header line has elements separated by “|”, where the sample name and locus name are the first and second elements, respectively. Any other elements after the sample and locus names, such as barcode, group, and read IDs, are ignored by the script. Four clustering identity thresholds specified by the `-c/--clustering_identities` flag and two size thresholds specified by the `-s/--size_threshold` function identically to their respective parameters in the config file.

The reclustering script produces output organized similarly to the `PURC v2.0` output detailed above. This output contains two FASTA files for each locus, one with the OTU sequences combined from all samples and the other containing an alignment of those sequences. There are folders for each locus, and within each, separate folders containing working files for each sample. A summary file called `purc_cluster counts.xls` contains information about number of reads, OTUs, and chimeras found per sample and per locus.

## 4 Conclusions

`PURC` (Rothfels et al., 2017), in conjunction with PacBio circular consensus sequencing, introduced an economical and effective alternative to time-consuming cloning and Sanger sequencing for generating broad multi-locus datasets for groups containing polyploids. With `PURC v2.0` we have significantly improved upon the earlier version, most notably by addressing shortcomings of OTU clustering by the incorporation of ASV inference. In addition to the generally greater accuracy of ASV (versus OTU) inference, our implementation of “straight” ASV inference and ASV inference with priors alongside OTU inference allows users to compare the results from all three approaches. `PURC v2.0` can thus function as a data exploration tool, providing users with the opportunity to detect and interrogate unexpected patterns of variation in their amplicon sequencing datasets.

We anticipate that `PURC v2.0` will prove to be a valuable component of biologists’ toolkits. While amplicon-based data generation does not scale as well as most other reduced-representation techniques, such as hybseq (*e.g.*, Breinholt et al., 2021; Johnson et al., 2019), it is a cost-effective way to sequence a greater number of samples for fewer—but more informative—loci, allows for the easy integration of new data with historical datasets, and is a powerful means of supporting “moderate data” approaches (*i.e.*, the production of multi-locus datasets that are large enough to be phylogenetically informative yet sufficiently small to allow for thorough curation and model selection; Rothfels et al., 2015, 2012, see also Frost and Lagomarsino, 2021). Moderate data are particularly effective for the phylogenetic study of polyploids, where the limiting factor is typically systematic error rather than stochastic error (Philippe et al., 2011): the accurate recovery and analysis of the full set of homeologous sequences (avoiding chimeras, etc) is more important than the absolute amount of data available *per se* (Rothfels, 2021).

The main application of `PURC v2.0` is thus likely to be as a component of a “polyploid phylogenetics” workflow (Rothfels, 2021). For example, a researcher can use `PURC v2.0` to generate a multi-locus nuclear dataset for a broad taxon sample, including polyploids, phase the loci (determine, for each locus, which copy of each polyploid comes from which subgenome) with homologizer (Freyman et al., 2020, Freyman and Rothfels, this volume), and use those phased multilocus data for downstream phylogenetic inferences such as divergence-time or species-tree estimation. Such a workflow would allow for the investigation of many outstanding questions related to polyploid evolution, and would also permit the phylogenetic study of groups that contain polyploids regardless of whether polyploidy itself is of central interest. In addition, `PURC v2.0` is not restricted to cases of the multiple copy problem. For example, to answer questions of hybrid parentage, plastid regions can be amplified and co-sequenced with nuclear loci and processed in the same `PURC` run, and `PURC v2.0` is also an effective way to generate sequence data for non-hybrid diploids. Finally, while not specifically designed for metabarcoding, the underlying programs in `PURC v2.0` are widely used in this field, making `PURC` useful for demultiplexing samples and sequence inference for other downstream analyses (*e.g.*, Goldberg et al., 2020).

## Notes

https://bitbucket.org/peter_schafran/purc/src/master/

### Competing Interest Statement

The authors have declared no competing interest.

